# Double haploid development in drug-type *Cannabis sativa* L. through microspore indirect *de-novo* plant regeneration

**DOI:** 10.1101/2024.10.25.620185

**Authors:** Francesco Tonolo

## Abstract

Double haploid technology (DH) is an essential tool in plant breeding, enabling the rapid production of homozygous lines. However, *Cannabis sativa* L. has been categorized as recalcitrant to DH induction. In this study, we evaluated the potential to generate DH *C. sativa* plants via anther culture and indirect *de-novo* organogenesis. We examined a THCA-dominant cultivar with a callus induction success of 29.48%. Mixoploidy in the callus indicated spontaneous genome doubling. This is the first report documenting the successful induction of DH *C. sativa* plants through *de-novo* indirect organogenesis. These findings have profound implications for the *C. sativa* breeding sector by potentially improving efficiency of genome editing and hybrid development in this economically significant species.

## Introduction

The plant kingdom is characterized by a high level of developmental plasticity and totipotency, which enable single cells other than the zygote to develop and differentiate into a mature plant (Su et al., 2021). Scientists exploit this natural survival strategy in several ways to study, engineer, manipulate, and breed plants. Indeed, the possibility of studying and working on a single cell that eventually becomes a fully developed plant is extremely advantageous.

There are two main examples of totipotency, somatic or gametophytic, each of which can take two different developmental routes: the embryogenesis or the *de-novo* organogenesis pathway (Soriano et al., 2013; Long et al., 2022). The main differences are determined by the type of cells that can proliferate and the developmental route which leads to a fully regenerated plant. The origin cells can be either gametes or somatic cells. At the same time, the developmental route can either involve the generation of an embryo or the differentiation of the meristematic center in different organs (Lardon & Geelen, 2020). In the case of somatic regeneration, the cells originate from a diploid vegetative tissue. The regenerated plant generally presents the same genetic profile and ploidy level as the donor plant, although this process can also contribute to generating plants with new characteristics due to somaclonal variation (Wang & Wang, 2012; Galán-Ávila et al., 2020).

On the other hand, gametophytic proliferation is a form of totipotency based on the proliferation of the male or female haploid gametes and the associated cells (Seguí-Simarro, 2010). In this case, the cells that proliferate are derived from meiosis, and therefore, they represent the haploid segregant progeny of the donor plant. Apart from having a unique genetic profile, these cells have a different ploidy level as they are generated by a haploid reproductive cell.

These ploidy levels and genetic profile changes have been used in several ways to advance the human understanding and exploitation of plants’ survival strategies. Indeed, once a haploid plant is generated and it undergoes genome doubling spontaneously or artificially, a so-called double haploid (DH) is obtained. Because of this process, the resulting DH plant is completely homozygous and obtained in just one generation (Castillo et al., 2009).

The double haploid plants have been exploited by scientists to develop immortalized molecular mapping populations, to fix traits obtained through genome editing techniques quickly, or to simplify genome sequencing by eliminating heterozygosity (Ferrie & Möllers, 2011). Moreover, DH technology has proven paramount for the breeding sector, given the quickness in the generation of homozygous lines for F1 hybrid production, the rapid fixing and introgression of new traits and the exploitation of the gametoclonal variation and *in-vitro* selection system to decrease time, labour, and costs of plant-breeding programs significantly (Germanà, 2011; Rajcan et al., 2011).

The DH technology has been applied to many plant species (Chen et al., 2020) with varying results. It has been documented that the successful generation of DH plants is species and genotype-dependent, with the need to fine-tune the inputs for successful outcomes. The low microspore proliferation, embryo formation, and plant regeneration frequency are challenging bottlenecks that scientists have to overcome. For these reasons, several species of agronomic interest, such as *Solanum lycopersicum, Solanum melongena*, and *Capsicum annuum*, have been deemed recalcitrant to the process (Seguí-Simarro et al., 2010).

Given the undoubted usefulness of this biotechnology, several efforts have been made to apply it to *Cannabis sativa* L. (Cannabaceae). Indeed, *C. sativa* is an allogamous plant species that has been widely cultivated due to its industrial (Karche & Singh, 2019), ornamental (Hesami et al., 2022), nutritional (Kruger et al., 2022), and particularly for its broad medicinal potential (Andre et al., 2016).

While most drug-type *C. sativa* breeding has historically happened underground, the legal drug-type *C. sativa* breeding sector currently has an estimated yearly value of 1.29 billion USD and a projected value of 8.08 billion USD for the year 2030 (Deore, 2023). Successful application of the DH technology on *C. sativa* would, therefore, be an innovative and extremely valuable biotechnological application.

To date, there are only a few reports on microspore embryogenesis induction in *C. sativa* aiming to develop DH plants. Both reports documented a successful induction but a failure to yield viable embryos able to progress further into their development (Adhikary et al., 2021; Galán-Ávila et al., 2021). Additionally, all the most relevant studies from 1972 up to 2020 evaluating the potential of *in-vitro* plant regeneration have been summarized by Adhikary et al. (2021). This review documented varying degrees of success of indirect (via callus culture) and direct (via meristematic tissue culture) plant regeneration. The data shows that direct *de-novo* organogenesis often appears to yield good results, while indirect regeneration via callus formation generally has regeneration rates that are absent or relatively low. The failures of the last 20 years and the challenges faced by the scientists, therefore, prompted the researchers to classify *C. sativa* as a recalcitrant species to plant regeneration and DH induction (Galán-Ávila et al., 2021; Monthony et al., 2021;).

Here, with several experiments, we set out to study the potential generation of DH *C. sativa* plants through anther culture and the regeneration of plants via gametophytic indirect *denovo* organogenesis.

The experiments were designed based on several crucial factors. First, given the long history of failures documented in the literature, the previous results were used as landmarks to give the DH program a fair shot. Indeed, the observations regarding the failure in the progression of the embryogenic pathway during gametophytic proliferation and the absence of reports on somatic embryogenesis would tell us that this might not be a viable route, although it is the preferred one. Moreover, while the microspore culture is the preferred method, it has been observed that in the case of recalcitrant species where microspore culture was unsuccessful, anther culture was the preferred route (Soriano et al., 2013).

Secondly, the experiments were designed to be broadly relevant by including multiple genotypes and selecting widespread commercial cultivars encompassing both *C. sativa* CBDA and THCA drug-type gene pools and including male and “masculinized” female plants.

Thirdly, to tackle the drawbacks of the anther culture, namely the uncertainty about whether the origin cells were gametes or somatic cells of the anther wall, we designed a ploidy and genotyping test to detect DH plants and, therefore, determine whether the obtained plants were indeed derived from microspores (Germanà, 2011).

With these experiments, we set out to answer the following questions:

1) Can we induce callus growth from *C. sativa* anther culture?
2) Can we induce indirect organogenesis in the callus? And can we regenerate a plant from callus?
3) What is the ploidy level of the callus and regenerated plant?
4) Is the obtained callus derived from the proliferation of a microspore?

## Material and methods

### Plant material and growth conditions

For the experiment 1 *C. sativa* seeds of a commercial THCA dominant drug-type cultivar Hash Plant were sourced. The cultivar was chosen for its high cannabinoid productivity and extensive use as a parental in breeding programs to produce commercial drug-type varieties. This choice would make results more broadly applicable to modern drug-type varieties (Fig S1). Plant growth was carried out following standard agricultural practices for producing *C. sativa* inflorescences, as previously described by Magagnini et al. (2018). Upon changing the photoperiod to short daylight conditions (12h/12h light/darkness), plants flowered, and anthers were collected after three weeks from male plants in experiment 1.

### Pollen microscopy

During experiment 1, pollen was isolated from anthers by dipping them in deionized water and gently squishing the anthers between the glass slides. The slides were checked under the microscope (Leica Microsystems GmbH, Wetzlar, Germany) using the differential interface contrast technique to assess their developmental stages and determine which anthers were most suitable for the callus induction process.

### Ploidy measurements

The method for ploidy analysis was carried out following a modified protocol by Zonneveld and van Iren (2001). Nuclei isolation buffer containing 4.15g/l MOPS, 9.15g/l MgCl2, 8.8g/l TriSodium citrate, 1.55g/l DTT, 25g/l Polyvinylpyrrolidone and 1ml/l Triton-x was prepared before the analysis and stored at 4°C. Subsequently, 100-200mg of fresh plant material was collected and kept on ice until analysis. The samples were prepared by chopping the plant material with a razor blade in a Petri dish containing 1ml of buffer. The buffer containing the free nuclei was then filtered through a 20μm Minisart nylon filter (Sartorius, Goettingen, Germany) into a 1.5ml Eppendorf and kept on ice while the other samples were processed. After all samples were prepared, Propidium iodide was added to a final 50μg/ml concentration. The samples were mixed and incubated in darkness on ice for 10 minutes before analysis. Ploidy analysis on a flow cytometer (Milliliter Guava Easycyte, Merck, Darmstadt, Germany) was performed following analytical parameters as stated by Parson et al. (2019) and Kurtz et al. (2020). The laser at 488nm was used for excitation, and fluorescence was recorded with the Yellow H linear channel at 583/26nm. Leaves of diploid *C. sativa* plants were used as standards. Samples were measured in triplicate.

## Results

### Experiment 1

Male plants of a THCA dominant cultivars were grown and anthers collected. Anthers were examined under differential interface contrast (DIC) microscopy. Imaging of the microspores revealed the presence of microspores at different developmental stages according to the development of the anthers (Fig 1). Indeed, tetrads were found in the smallest sampled anthers, newly released microspore and polarized microspores were found in medium size anthers and developed microspore were found in the biggest sampled anthers (Fig 2).

**Fig 1.**
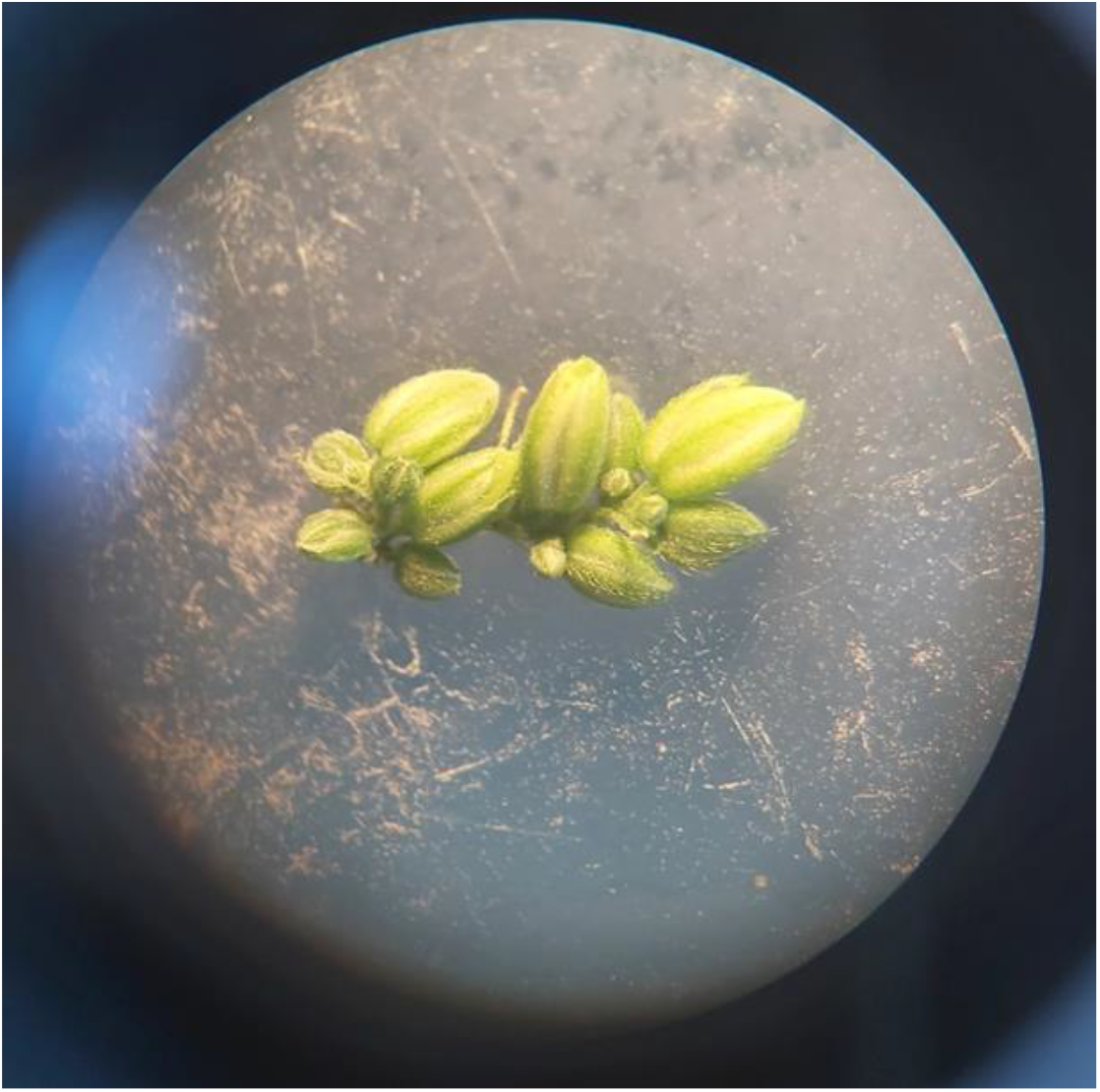
Anthers at different developmental stages observed at the stereo microscope.

**Fig 2.**
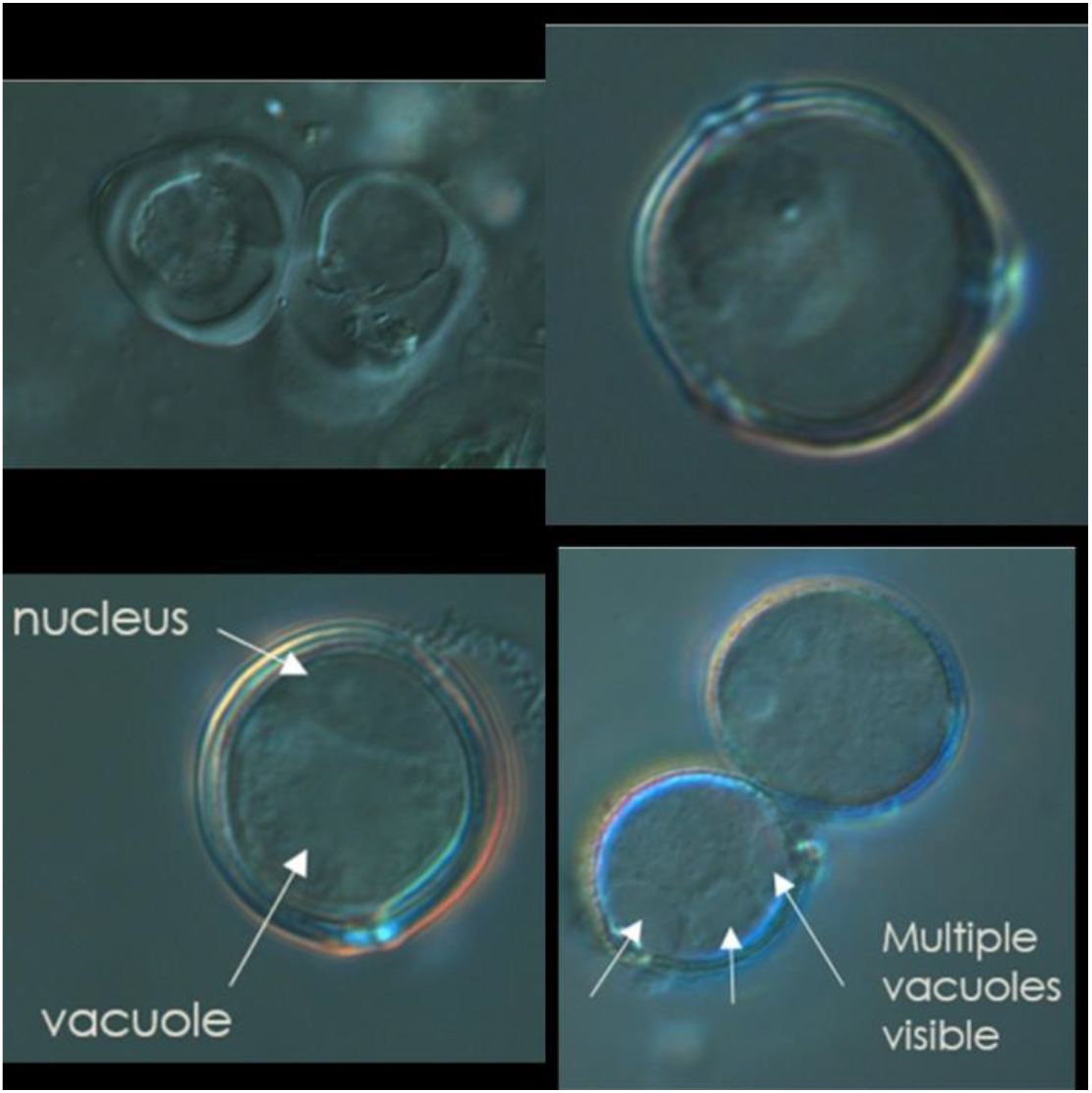
DIC microscopy imaging of pollen in different developmental stages. In clockwise order from top left panel: tetrad, newly released microspore, developed microspore, polarized microspore.

Anthers were selected, surfaced sterilized and placed in callus induction media. After 4 weeks callus induction rate was assessed (Fig 3). Results showed an induction rate of 28.84% for genotype HP2, 23.91% for genotype HP5, 18.18% for genotype HP6 and 39.62% for genotype HP7. In total for the THCA cultivar under examination callus induction rate was 29.48% (Table 1).

**Fig 3.**
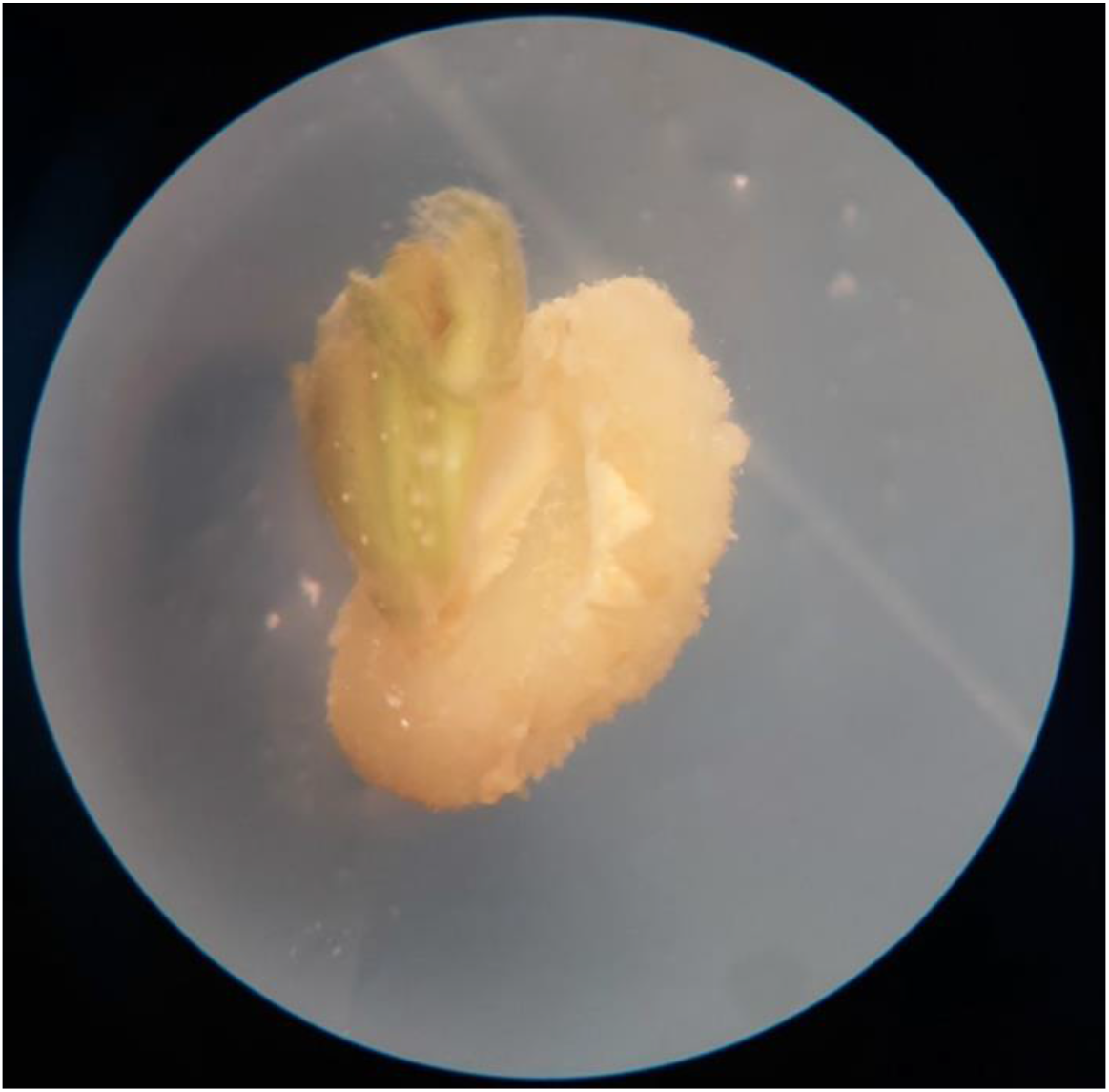
Callus growing out of the anther observed under the stereomicroscope after four weeks from inoculation during experiment 1.

**Table 1.** Results of the first experiment on the high THCA cultivar Hash Plant. NA stand for not assessed (contaminated plate). Results of root organogenesis are based on the biggest callus sample of each genotype chopped up to generate many smaller calli. The ploidy analysis was destructive therefore the two tested calli could not be used for the organogenesis assessment.

| Plant ID | Callus/anthers | Root organogenesis (one callus chopped up) | Shoot organogenesis/callus |
| --- | --- | --- | --- |
| HP2 | 15/52 (28.84%) | yes | 2/12 (16.67%) |
| HP5 | 11/46 (23.91%) | yes | NA |
| HP6 | 4/22 (18.18%) | yes | NA |
| HP7 | 21/53 (39.62%) | yes | 4/18 (22.22%) |
| Total for cultivar | 51/173 (29.48%) | yes | 6/30 (20%) |

The ploidy level of the callus was assessed by sampling two calli originating from each of the genotypes HP2 and HP7. Flow cytometric results showed that the callus was mixoploid in all samples tested (Fig 4). Interestingly, the callus did not present haploid cells (n) as it would be expected from the anther culture. On the other hand, diploid (2n), tetraploid (4n), octoploid cells (8n), 16n and 32n cells were detected.

**Fig 4.**
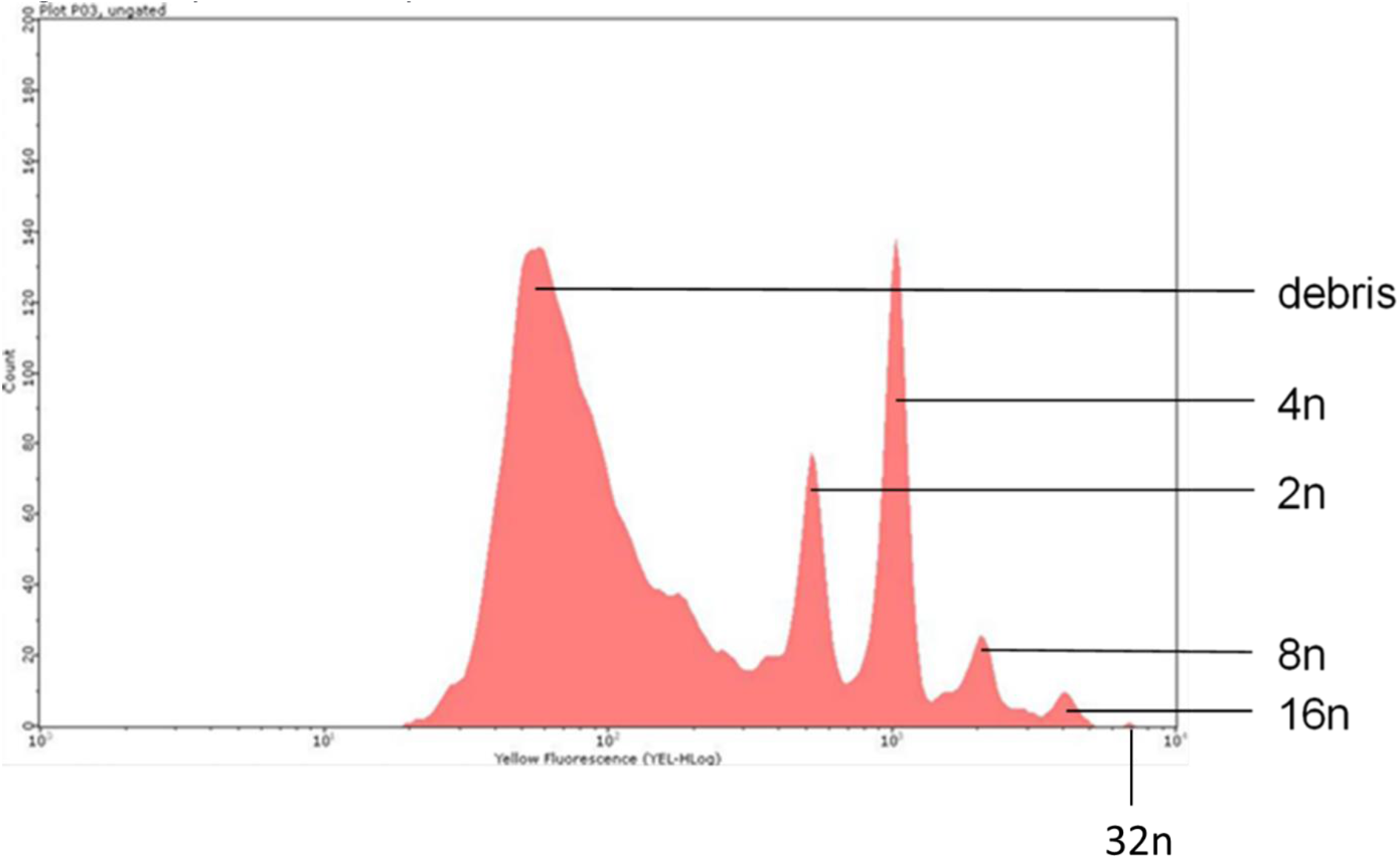
Flow cytometric analysis of callus obtained from anthers culture of HP2.

To assess the capacity of indirect root organogenesis the biggest callus mass was selected from each of the genotype under study. These calli were cut in many smaller pieces and placed in root induction liquid media (Fig 5A). This resulted in successful root induction for all 4 genotypes, with ample production of roots within four weeks of inoculation. Assessment of indirect shoot regeneration was instead evaluated on the remaining calli of all four genotypes. Calli were plated on shoot induction solid media and evaluated after 4 weeks (Fig 5B). Unfortunately, the plates of HP5 and HP6 were contaminated and therefore discarded. For the genotypes HP2 and HP7 the shoot induction success was 16.67% and 22.22%, respectively. On average shoot organogenesis success for the cultivar under study was 20% (Table 1).

**Fig 5.**
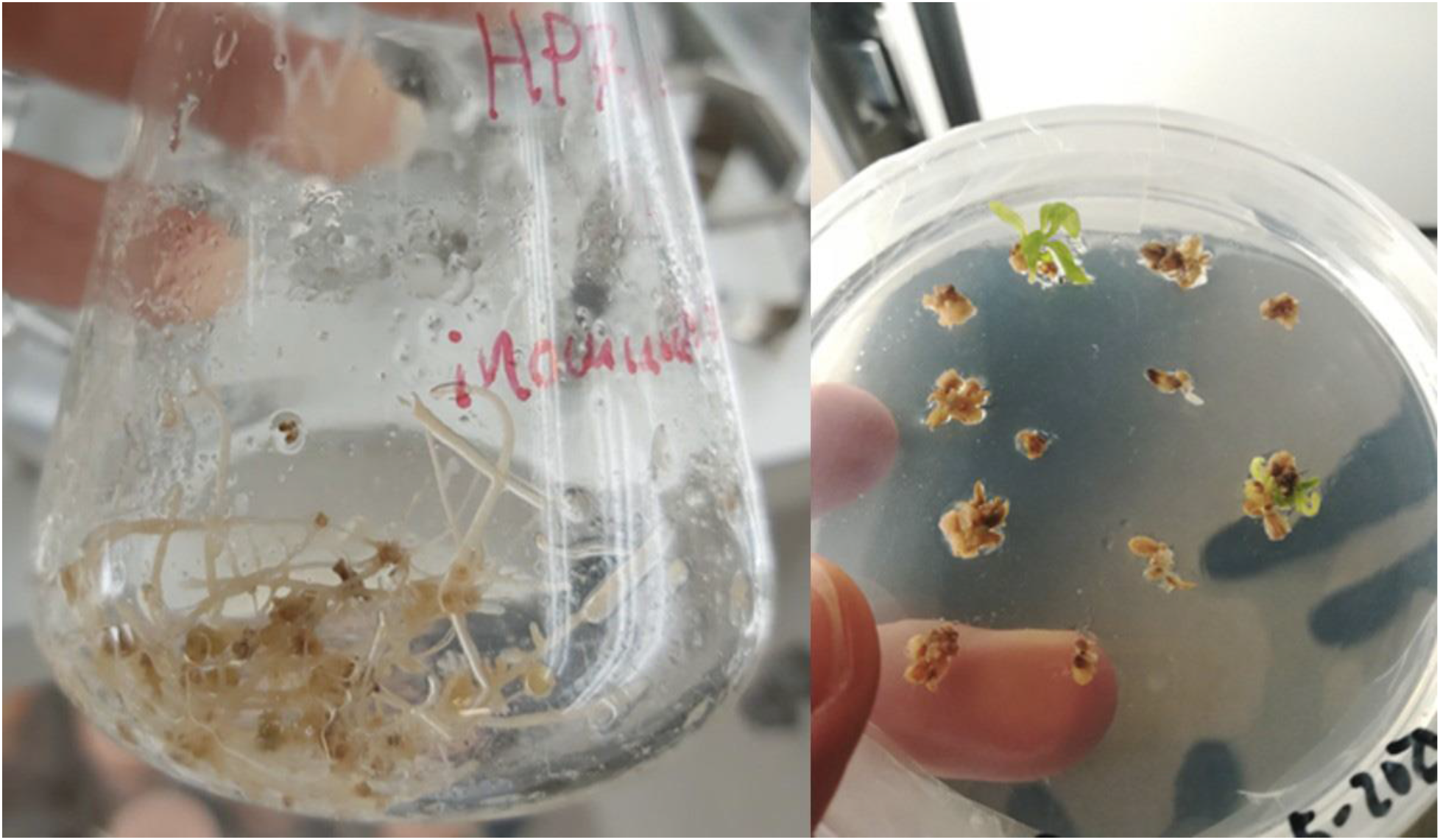
Regenerated roots of genotype HP7 (**A**) and shoots of genotype HP2 (**B**) from callus during experiment 1.

### Experiment 2 (data not shown to avoid potential disputes)

## Discussion

The results of the first experiment revealed several significant findings.

First, the microspore imaging revealed that the anthers contained microspores at all four stages of development. Interestingly, there was a correlation between the size of the anthers and the levels of microspore development. These observations agreed with Gálán-Álvida et al. (2021), who also found a correlation between the dimension and advancement in the development of the anthers and the microspores they contained.

Secondly, the induction of callus was observed for all four genotypes tested with an average of 29.48% over the cultivar, demonstrating that regardless of the genotype under study, they all produced callus at a relatively good rate. Moreover, the same was true for the organogenesis process. The induction of roots was highly successful for all genotypes, with ample production of healthy-looking root structures. In the case of shoot organogenesis, although two genotypes were lost due to contamination, we still could evaluate the capacity of the cultivar Hash Plant to regenerate shoots based on the remaining two genotypes. The results showed that shoot organogenesis could be obtained from both genotypes with an average of 20% over the cultivar.

Thirdly, the ploidy analysis revealed several unexpected findings. The expectation of this test would be that being the microspore haploid (n) if these gametophytic cells would proliferate and grow into a callus, we should find a haploid callus, yet this was not the case. On the other hand, if the cells originated from diploid somatic cells of the anther wall, we should observe a diploid callus, yet this was not the case. Interestingly, we detected a mixoploid callus, which presented cells at several ploidy levels: diploid (2n), tetraploid (4n), octoploid (8n), 16n cells and even 32n.

While the 32n cells can be explained as a population of 16n cells during the S and G2 phases of the cell cycle, this does not explain the cells at the lower ploidy levels.

In this regard, it has been documented that haploid genomes in plants are subjected to instability (Germanà, 2011). In order to evade this genomic state, cells can double up their genome to stabilize their genetic structure and to survive the stressful *in-vitro* culture conditions under undifferentiated growth. In many crops, processes like endomitosis, nuclear fusion, or endoreduplication result in spontaneous genomic duplication during *in-vitro* culture (Seguí-Simarro & Nuez, 2008).

Endoreduplication has been documented in various plant species, is particularly common in reproductive tissues and is often associated with cell growth, differentiation, and stress response (Leitch & Dodsworth, 2017; Lang & Schnittger, 2020).

The mechanism at the base of this process involves changes in the regulation and abundance of a variety of cyclin-dependent kinases, cyclins and regulatory proteins/transcription factors. This results in cells skipping the M phase while cycling between the G and S phases of the cell cycle (Fig 13).

**Fig 13.**
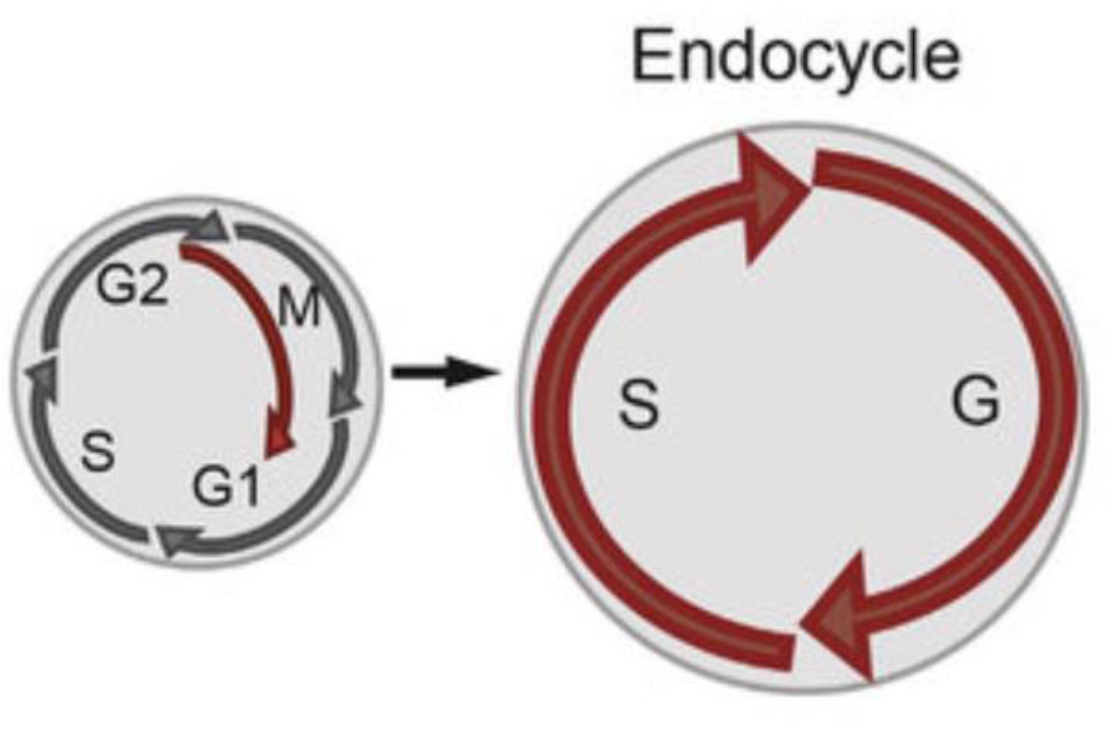
cell cycle during endoreduplication

The fact that 2n, 4n, 8n, and 16n cells were observed suggests that this process occurs in *C. sativa* callus when growing *in-vitro*. It appears that the microspores with their haploid genome might not be able to survive during the *in-vitro* culture and necessitate doubling up their genome. The fact that we cannot detect haploid cells also suggests that this process occurs very early during the culture and that only cells of higher ploidy levels can develop. These findings agree with the observations of Haplotech Inc. (Adhikary et al., 2021) and Galán-Ávila et al. (2021), which observed the death of *C. sativa* haploid embryos early on during their development.

Moreover, the fact that we detected 4n, 8n, and 16n cells supports this hypothesis, as genome doubling occurred at least four other times to give rise to these higher-ploidy cell populations.

Overall, it appears that the haploid microspores can divert their developmental fate by generating callus early during their proliferation. At the same time, the undifferentiated callus growth elicits repetitive genome doubling events, which are likely necessary for survival.

Indeed, this culture system provides an incredibly large number of diverse microspores derived from the reshuffling of the genomic information of the donor plant’s parents and extreme artificial selection pressure. Essentially, this *in-vitro* system replicates and takes advantage of natural biological systems geared toward the selection of the fittest by exploiting the genetic diversity of microspores, their totipotent characteristics, and the capability of ploidy level jumps through endoreduplication and mitosis to regenerate fully homozygous diploid *C. sativa* plants from haploid cells.

## Conclusion

In conclusion, with these experiments, we could successfully induce callus growth from *C. sativa* anther culture, induce indirect *de-novo* shoot and root organogenesis from the obtained callus, and ultimately regenerate and acclimatize several plants with this system. The investigations based on the ploidy tests could tell us more about the underlying processes occurring during the *in-vitro C. sativa* culture. Indeed, while the ploidy measurements did not reveal haploid cells, the results were fundamental for documenting and understanding the shifts of ploidy levels necessary for this process to be successful. On the other hand, the genetic test was fundamental to confirm the DH nature of the obtained plants.

To the best of our knowledge, this is the first report on the successful induction of double haploids in *C. sativa* leveraging protocols that can regenerate plants via the indirect *de-novo* organogenesis pathway.

Overall, the designed culture system has several advantages, making it an extremely valuable asset for the *C. sativa* breeding sector.

Firstly, given the allogamous nature of this species, inbreeding depression and deleterious mutations are a real threat to the progression of breeding programs. Indeed, a loss of vigor and fertility is often observed upon full-sib or selfing inbreeding (Carlson et al., 2021).

This culture system provides a solution as it is designed to select microspores presenting a genetic profile suitable for prolific growth and their ability to regenerate into a plant, essentially developing homozygous lines in one generation while at the same time purging recessive deleterious alleles, making this technology paramount for the development of F1 seed lines.

Moreover, efficient indirect plant regeneration is paramount for the successful application of biotechnologies, which have many bottlenecks with low success rates, such as genetic transformation with *Agrobacterium* and CRISPR-Cas systems (Zhang et al., 2021). With its high regeneration efficiency, the designed culture system is the perfect base for implementing genome editing biotechnologies with low-efficiency rates. Indeed, the haploid microspore is the most desired target for genetic transformation using CRISPR or other genome-editing DNA-free systems (Bhowmik et al., 2018). When the mutations occur in the haploid genome and then the mutated cell, through the process of regeneration, reaches the diploid status, it results in a homozygote mutation. Thus, with this process, we can potentially avoid chimeric plants and obtain a fully expressed mutation without further crossings, speeding up the lab work and breeding process significantly. Therefore, this culture system is a gateway to unlock the potential of modern genome editing techniques on *C. sativa* (Adhikary et al., 2021), enabling the development of new cultivars at a quicker pace and more cost-effectively while at the same time providing the much-needed genetically healthy and stable starting material for F1 breeding pipelines.

---

**Fig S1.**
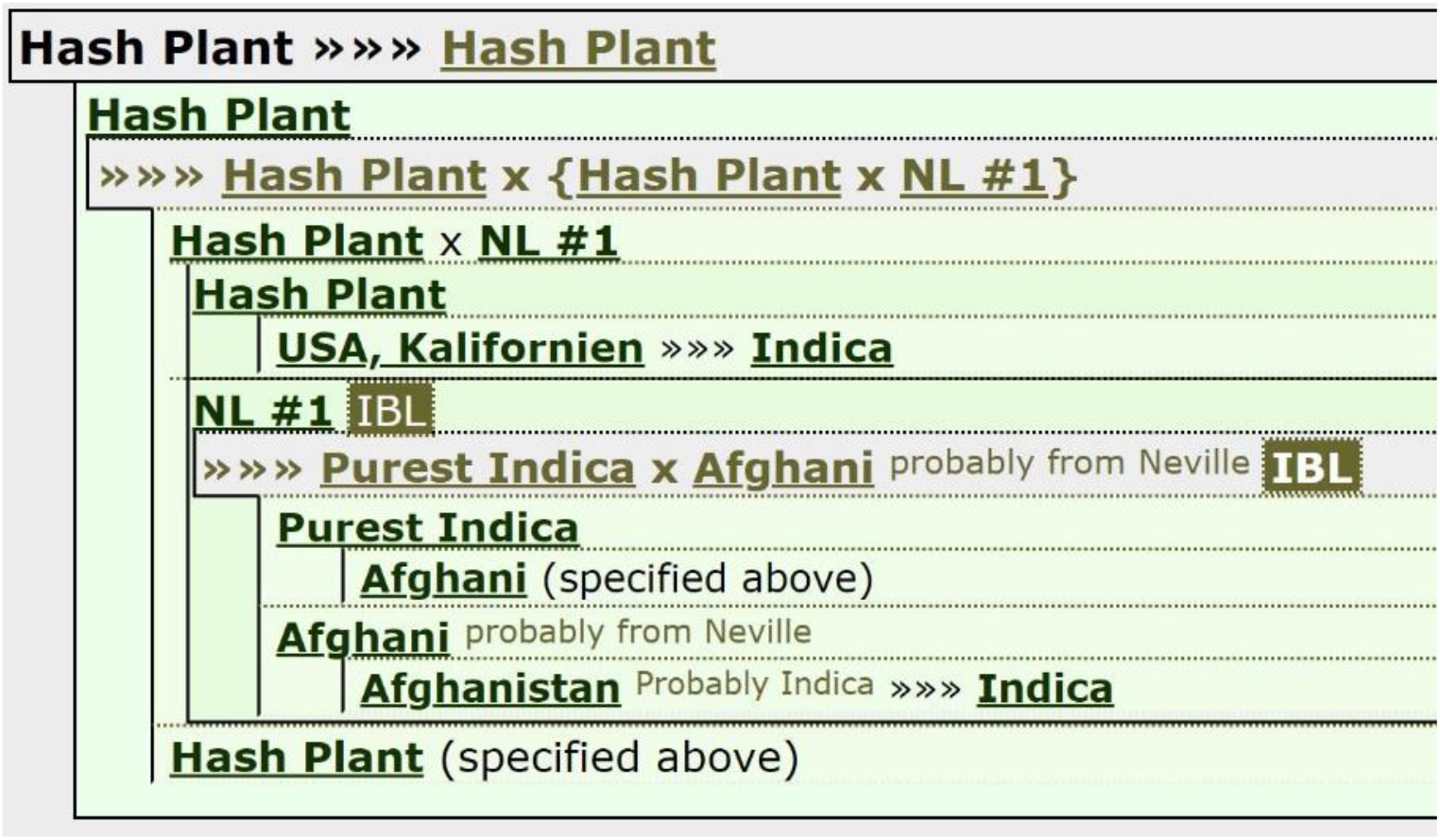
Genetic background of Hash Plant used in experiment 1

